# Streptococcus suis infection on European farms is associated with an altered tonsil microbiome and resistome

**DOI:** 10.1101/2022.08.01.500980

**Authors:** Simen Fredriksen, Carlos Neila-Ibáñez, Isabel Hennig-Pauka, Xiaonan Guan, Jenelle Dunkelberger, Isabela Fernandes de Oliveira, Maria Laura Ferrando, Florencia Correa-Fiz, Virginia Aragon, Jos Boekhorst, Peter van Baarlen, Jerry M. Wells

## Abstract

*Streptococcus suis* is a Gram-positive opportunistic pathogen causing systemic disease in piglets around weaning age. The factors predisposing to disease are not known. We hypothesised that the tonsillar microbiota might influence disease risk via colonisation resistance and/or co-infections. We conducted a cross-sectional case-control study within outbreak farms complemented by selective longitudinal sampling and comparison with control farms without disease occurrence. We found a small but significant difference in tonsil microbiota composition between case and control piglets (n=45+45). Variants of putative commensal taxa, including *Rothia nasimurium*, were reduced in abundance in case piglets compared to asymptomatic controls. Case piglets had higher relative abundances of *Fusobacterium gastrosuis, Bacteroides heparinolyticus*, and uncultured *Prevotella* and *Alloprevotella* species. Despite case-control pairs receiving equal antimicrobial treatment, case piglets had higher abundance of antimicrobial resistance genes (ARGs) conferring resistance to antimicrobial classes used to treat *S. suis*. This might be an adaption of disease-associated strains to frequent antimicrobial treatment.

## Introduction

*Streptococcus suis* is a Gram-positive bacterium that colonises the upper respiratory tract of pigs. It is one of the major bacterial causes of disease in pigs and a zoonotic pathogen causing sepsis and meningitis in humans [1–4]. Swine infections are prevented by metaphylactic use of antimicrobials. This has led to increased antimicrobial resistance (AMR) in *S. suis* isolates [5, 6], with macrolide and tetracycline resistance genes *erm(B)* and *tet(O)* being the most common [7]. The spread of antimicrobial resistance genes (ARGs) in zoonotic *S. suis* and to other streptococci is of concern to veterinary and human medicine [8].

The palatine tonsils have been suggested as a main habitat for *S. suis* colonisation and a site of entry into the host bloodstream [9–17]. A recent study identified differences in taxonomic composition of the tonsillar microbiota in piglets diagnosed with *S. suis* disease [18]. Microbiota associations with *S. suis* and other infectious diseases are of great interest to understand co-infection dynamics and to identify candidate probiotic species that may confer colonization resistance [19]. It is thought that co-infection with bacterial and viral pathogens of the porcine respiratory disease complex (PRDC) may predispose to *S. suis* invasive disease [20–22].

Tonsillar colonisation by different *S. suis* strains may also have an impact on invasive disease risk [14, 23]. While *S. suis* has been described as an opportunistic pathogen, different strains have varying virulence potential and can be grouped into commensal and pathogenic clades [24–28]. Strains from pathogenic *S. suis* clades have reduced genome sizes [25] and putatively reduced ability to persist as colonisers in competition with commensals. Strains from pathogenic clades (i.e., disease-associated strains) can also be isolated from the tonsils of healthy pigs, but the prevalence of asymptomatic carriage of disease-associated strains, and whether invasive *S. suis* disease is preceded by outgrowth of these strains, is not known.

The aim of this study was to identify the tonsillar microbiota associated with *S. suis* systemic disease in piglets at weaning age. We utilized a case-control study design and next-generation sequencing to characterise the tonsillar microbiota composition of piglets with systemic *S. suis* infection and co-housed asymptomatic piglets from nine European farms and compared these to piglets from four farms without *S. suis* disease. We used 16S rRNA gene amplicon- and metagenomic shotgun sequencing to quantify the tonsillar microbiota and resistome, and analysed clinical and non-clinical *S. suis* strains from the farms by whole-genome sequencing. This study design allowed us to assess microbiota composition and ARG abundance, as well as to assess the abundance of virulence marker genes prevalent in clinical or non-clinical *S. suis* strains.

## Results

### Sequencing

We sampled 45 case-control pairs of piglets with clinical signs of systemic *S. suis* disease and asymptomatic control piglets from the same pen. Case-control pairs were matched for age, genetic background, diet, and antimicrobial treatment. Additional control piglets from case farms and from control farms without *S. suis* cases were sampled to assess differences between farms. We also sampled one US farm and naturally weaned piglets raised organically in a forest for comparison (see Supplementary file 1 for farm information). Amplicon sequencing of 295 samples yielded an average of 76757 reads per sample after processing with DADA2 [29]. Metagenomic shotgun sequencing of 109 samples, including all case-control pairs and a subset of piglets from control farms, yielded an average of 7.5 million reads per sample after removal of host DNA.

### The *Streptococcus suis* disease-associated microbiota

Case and control piglets on outbreak farms had significantly different tonsillar microbiota composition, but the effect size was small (R2 = 0.01 and p = 0.02, PERMANOVA on Bray-Curtis dissimilarity comparing case and control piglets within outbreak farms). Analysis per country showed varying case-control results for Spain (R2 = 0.030, p = 0.01) and Germany (R2 = 0.017, p = 0.93), which had fewer samples from more farms. The largest difference in tonsil microbiota composition was found on the Dutch farm, NL1 (R2 = 0.26, p < 0.01), but this may be because all 3 case piglets from this farm were sampled shortly after death, allowing opportunistic pathogens to bloom. However, we did not see a general association between the microbiota and clinical sign severity or (future) death within the other farms. There was no consistent difference in alpha diversity between case and control piglets (Figure S1).

We identified several amplicon sequence variants (ASVs) significantly associated with piglet case-control status within outbreak farms (Figure 1A). SIAMCAT [30] analysis identified three ASVs with significantly (p < 0.05) higher abundance in case piglets: ASV 29 (*Prevotella* sp., mean abundance: case: 1.8%, control: 0.7%), ASV 3 (*Alloprevotella* sp., case: 3.6%, control: 2.2%) and ASV 21 (*Fusobacterium gastrosuis*, case: 3.6%, control: 1.5%). These taxa were all part of the core microbiota (i.e., ASVs with at least 99% prevalence in control piglets) in the European farms. The case-associations of these three taxa were stronger in the Dutch and Spanish farms, where the outbreaks were larger and/or the symptoms more severe than in the German farms. These 3 ASVs also had the strongest association with case piglets using Wilcoxon rank sum test (p < 0.005), but no ASVs were significantly different when multiple testing correction was applied (FDR > 0.05). Furthermore, no genera were significantly different in abundance between case and control piglets (FDR > 0.05). Table S1 shows the case-control associations for all ASVs and genera.

While the three case-associated ASVs had the strongest case-control associations, few other ASVs trended towards case-associations (Table S1). A larger number of different ASVs had weaker, non-significant (FDR > 0.05), control-associations. Among the core microbiota members, ASV 48 (*Clostridium disporicum*, case: 0.38%, control: 0.74%), ASV 36 (*Rothia nasimurium*, case: 0.47%, control: 0.91%) and ASV 6 (*S. suis*, case: 1.98%, control: 2.62%) had the strongest control associations. Among the non-core ASVs, the strongest control-associations were ASV 11 (*Actinobacillus minor*, case: 0.27%, control: 0.63%, not present in farms ES4 and NL1) and ASV 81 (*Terrisporobacter mayombei*, case: 0.22%, control: 0.40%). In terms of total relative abundance, ASV 1 (*Moraxella porci* /*pluranimalium*, case: 3.5%, control: 8.1%) showed the greatest difference between case and control piglets. ASV 1 had strong control associations on farm ES2 (case: 0.9%, control: 5.6%, being completely absent in several case piglets) and ES3 (case: 15%, control: 24%), but was absent in farm ES4 and NL1 and equally abundant in German case and control piglets.

**Figure 1:**
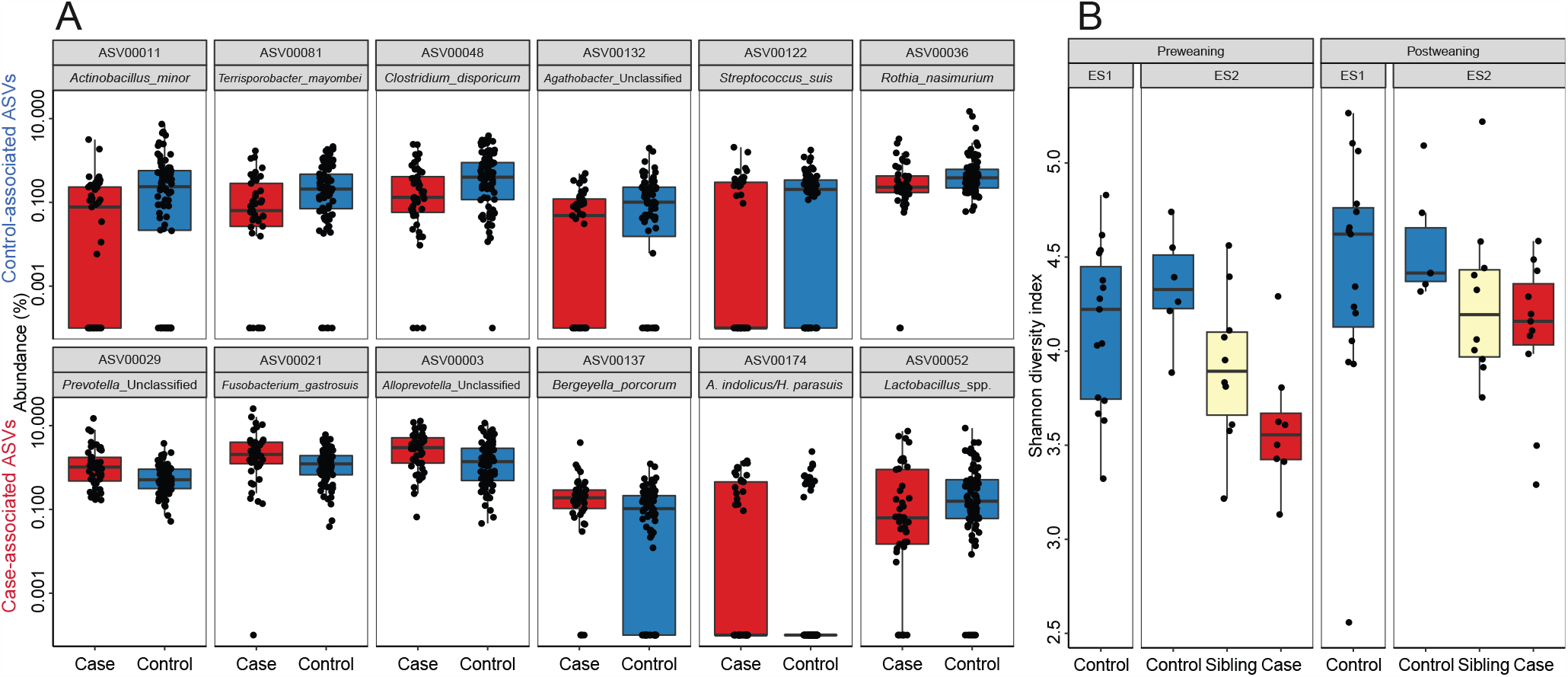
Microbiota case-control associations (amplicon sequencing). **A**. ASVs with the strongest unpaired Wilcoxon rank sum test case-control associations within the outbreak farms. Only ASVs with >0.1% mean abundance were included in this analysis. See Table S1 for statistics. Top row: control-associated ASVs, bottom row: case-associated ASVs. **B**. Shannon diversity of prospectively collected samples from pre-weaning piglets that developed *S. suis* clinical signs 2-4 weeks later, after weaning. Samples from future case piglets had lower Shannon diversity than control piglets that remained asymptomatic. Piglets from control farm ES1, which was managed by the same company and using sows from ES2, also had high mean diversity.

The porcine respiratory disease complex (PRDC) includes several viruses and bacteria that are thought to co-infect with *S. suis*. While PRDC is mainly associated with respiratory disease in older pigs, and this this study investigates weaning age systemic disease, we considered PRDC-associated bacteria to be of interest in possible co-infections. *S. suis* itself and *Glaesserella parasuis* were among the most abundant tonsil microbiota members. The relative abundance of *S. suis* ASVs trended towards being higher in healthy controls, whereas *G. parasuis* ASVs showed mixed associations. ASV 174 (100% identical to both *G. parasuis* and *Actinobacillus indolicus*) trended towards case-association (case: 0.27%, control: 0.16%). *Pasteurella multocida* was relatively prevalent (50%) but at low abundance (mean 0.06%), although two outlier case piglets had over 1% abundance. *Actinobacillus pleuropneumoniae* had 34% prevalence and 0.08% mean abundance but was most abundant in control piglets. *Bordetella bronchiseptica* had lower prevalence (17%) but higher abundance (0.10%) as some control piglets had up to 12% abundance. *Mycoplasma hyopneumoniae* was virtually absent from the dataset, with only 6 reads from a single control piglet.

### Early life microbiota diversity predicts onset of clinical signs weeks later

To assess the value of tonsillar microbiota composition in predicting future invasive *S. suis* disease, we collected prospective samples on farm ES2, which had a history of post-weaning *S. suis* outbreaks. We also collected samples at the same timepoints on farm ES1, a control farm without recorded *S. suis* problems despite using sows produced on farm ES2. We sampled a cohort of 40 piglets on farm ES2 one week before weaning, prior to visible clinical signs. Eight of the 40 piglets developed *S. suis* clinical signs 1-3 weeks post-weaning, and tonsil samples collected from these piglets post-weaning were included in the 45 case-control pairs.

Comparison of prospective samples collected from case and control piglets one week before weaning, i.e., 2-4 weeks before disease onset, showed that the tonsil microbiota of case piglets had significantly lower Shannon diversity compared to control piglets that remained asymptomatic (p = 0.005; Figure 1B). Case piglets trended towards having lower diversity also post-weaning, during the outbreak, but the effect was smaller and less significant (p = 0.12). Asymptomatic siblings of case piglets also had a lower diversity than control piglets from litters without case piglets and piglets from control farm ES1 (Figure 1B). However, the difference in composition was smaller pre-weaning (R2 = 0.06, p = 0.15, PERMANOVA on Bray-Curtis dissimilarity) than during the outbreak (R2 = 0.13, p < 0.01). The Spearman rank correlation coefficient between ASV ratio of case-control abundance pre- and post-weaning on the farm was low (R = 0.09, p = 0.4), indicating that separate ASVs were differentially abundant and driving case-control compositional differences pre- and post-weaning. The strongest pre-weaning prospective case-associations were ASV 88 (*Pasteurellaceae*, case: 1.4%, control: 0.55%) and ASV 186 (*Actinobacillus indolicus*/*minor*, case: 1.1%, control: 0.21%), while ASV 77 (Leptotrichia, case: 0.11%, control: 0.33%) and ASV 22 (Actinobacillus minor, case: 0.06%, control: 0.20%) had the strongest control associations (all p < 0.03, but none were FDR significant).

### Metagenome-assembled genomes (MAG) analysis

Shotgun sequencing was performed on samples obtained from all case-control pairs and a subset of piglets from control farms. Co-assembly per farm yielded 802 metagenome-assembled genomes (MAGs) (>70% completeness and <10% contamination). Read mapping to the MAGs revealed an overall case-control compositional difference similar to that found by amplicon sequencing analysis (Figure 2, R = 0.03, p < 0.01, Bray-Curtis dissimilarity PERMANOVA). Case-control associated MAGs largely corresponded to the ASVs identified by amplicon-based analysis, but some taxa, including case-associated *Bacteroides heparinolyticus*, were only identified by MAGs. The MAG approach resulted in stronger case-control associations (FDR < 0.0001) than the ASV approach, suggesting that the case-associated ASVs represented multiple strains with variable case-control associations. The two ASVs with the strongest case-association likely corresponded to a collection of different *Bacteroidales* MAGs. The MAGs with the strongest case-associations were classified as *Bacteroidales* UBA1309 and *Alloprevotella* F0040, clades poorly described and with few or no publicly available genomes from related isolates. Case-control associations of all MAGs are listed in Table S2.

**Figure 2:**
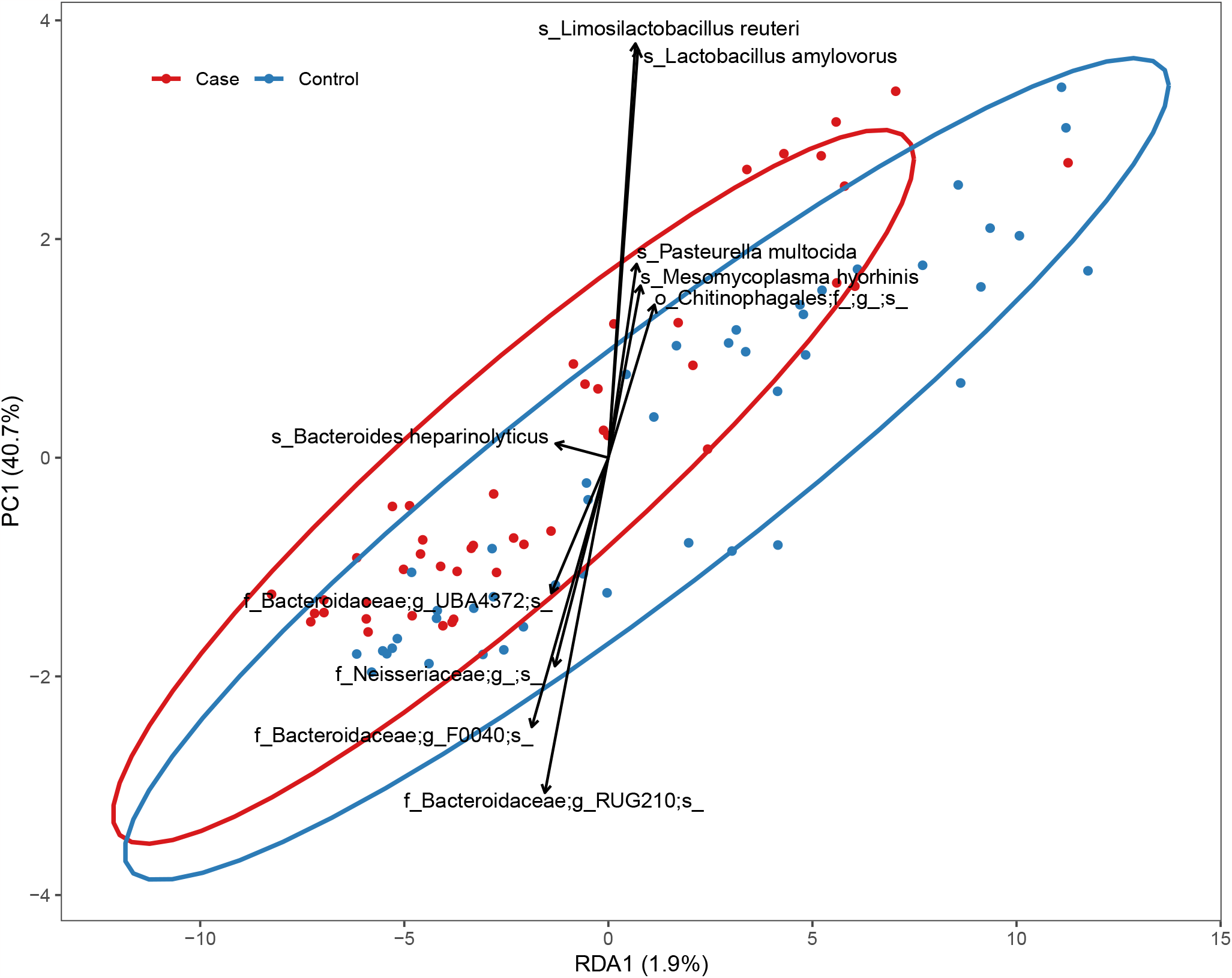
Redundancy analysis (RDA) on the abundance of metagenome-assembled genomes (MAGs) constrained by case-control status. The points represent individual samples. Arrows are response variables pointing in the direction of steepest increase of the values for the corresponding MAG; samples in the direction of the arrows have higher abundance of that taxon. Ellipses represent 95% confidence level.

### The tonsillar microbiome of piglets is similar between farms but diverged in free-range forest piglets

The tonsillar microbiota composition was largely shared between piglets within farms and countries (Figure 3A-B). Compared to the tonsillar microbiota of farm piglets, Tamworth free-range piglets kept in a forest in the Netherlands had a strikingly different microbiota and resistome (Figure 3C-D). While the European farm piglets shared a large core microbiota with 83 ASVs being present in 80% of piglets or more of the piglets, 44 of these were not found in the free-range piglets. Conversely, of the 157 ASVs found in all 5 forest piglets, 117 were not found in any farm piglet. The free-range piglets had low abundance of the genera most abundant in farm piglets, especially *Moraxella* (forest: 0.6%, farm: 13%) and *Streptococcus* (forest: 0.9%, farm: 12%), and higher abundance of *Acinetobacter* (forest: 10%, farm: 2.3%), “ *Rikenellaceae* RC9 gut group” (forest: 9.9%, farm: 0.1%) and *Treponema pedis* (forest: 3.6%, farm: 0.04%). The sample with the lowest *S. suis* abundance in the study was from a free-range piglet, which had 0.3% abundance of a single putative *S. suis* ASV (ASV 1050). This ASV did not have 100% identity to the 16S rRNA gene V3-V4 region of any *S. suis* strain publicly available in the SILVA or NCBI assembly databases. The other free-range piglets were colonised by *S. suis* ASVs shared with farm piglets.

**Figure 3:**
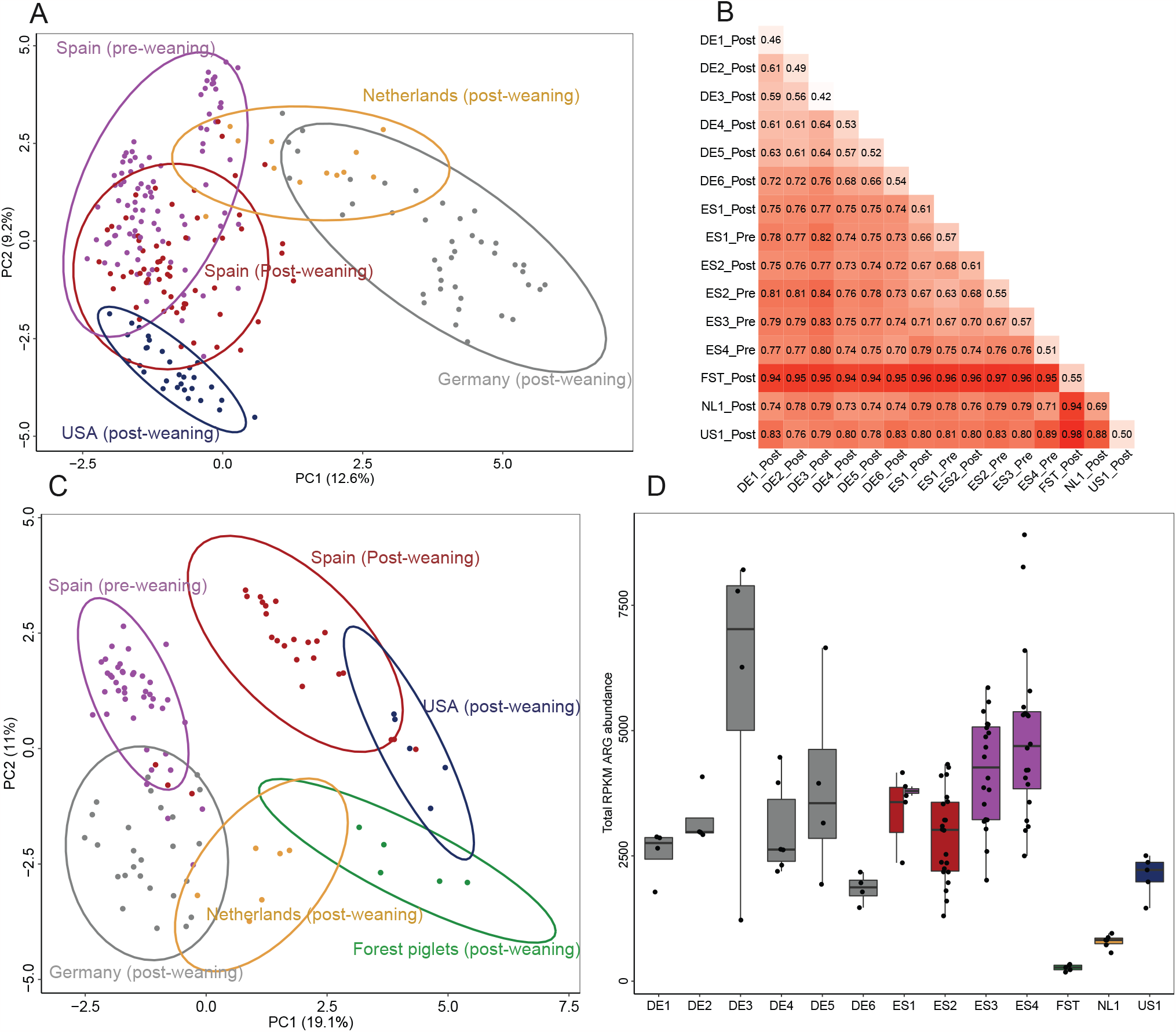
Microbiome differences between farms. **A**. PCA on microbiota composition (ASV abundance) of all pre- and post-weaning farm samples (excluding free-range piglets). The abundance was transformed with log(1000*abundance+1). Samples clustered broadly by country, but samples from farm DE6 clustered with NL1, and samples from ES4 clustered away from the other Spanish farms. Ellipses represent 95% confidence level. **B**. Mean pairwise Bray-Curtis dissimilarity between samples from the different farms. **C**. PCA on ARG abundance (transformed with log(1000*abundance+1)). The Spanish pre- and post-weaning samples separated into two separate clusters. **D**. Total abundance of all ARGs per farm.

### Increased abundance of tetracycline ARGs in case piglets

We quantified the abundance of antibiotic resistance genes (ARGs), collectively called the resistome, by mapping metagenomic reads to the Resfinder database [31] and normalising abundance by reads per kilobase reference per million fragments (RPKM). Piglets on most farms received antibiotics via feed or water, and these were included for resistome analysis. Twenty-two piglets received intramuscular injections, and these were excluded from the main resistome analysis. Each case-control pair included in the main analysis received equal antimicrobial treatment. Farms varied in total ARG abundance (Figure 3D), but shared high abundance of the most abundant ARGs, such as *blaROB-1*, which confers resistance to penicillin/amoxicillin/ampicillin (411 RPKM mean abundance), *sul2*, which confers sulfamethoxazole resistance (338 RPKM), and streptomycin resistance genes *aph(3’’)-Ib* (312 RPKM) and *aph(6)-Id* (220 RPKM). Table S3 lists all detected ARGs per farm. The most common ARGs in *S. suis, erm(B)* and *tet(O)* [7], were less abundant (66 and 111 RPKM, respectively). When comparing overall ARG composition by PCA, samples clustered by country, except for Spanish pre- and post-weaning samples, which clustered separately by age (Figure 3C).

In addition to taxonomic composition, ARG abundance may be influenced by both historical antimicrobial use on the farm and by ongoing antimicrobial treatment of the sampled piglets. We could not disentangle these two effects since most antimicrobial treatments were given equally to all sampled piglets on each farm. Farm NL1, a high health status research farm where the sampled piglets were not treated, and where piglets are rarely treated with antimicrobials, had the lowest ARG abundance (except for the free-range forest piglets). In Germany, the high health status farms DE1 and DE2 had low ARG abundance, but so did DE6, despite a history of severe *S. suis* disease. Farms DE3, ES3, and ES4 were assessed by veterinarians to have the lowest health status and highest historical antimicrobial use (Supplementary file 1), and piglets on these farms were also administered antimicrobials before sampling. These three farms had the highest ARG abundance (Figure 3D). There was no consistent link between the antimicrobials administered and the abundance of ARGs conferring resistance to these. On farm DE1, piglets that had received tetracycline had lower tetracycline ARG abundance than untreated piglets (810 vs 391 RPKM). On farm DE3, where all piglets had received tetracycline, tetracycline ARG abundance was lower than on DE2 and DE5 where piglets had received no antimicrobial treatment (804 vs 805 and 926 RPKM, respectively).

The total abundance of ARGs was 15% higher in case piglets compared to controls (case: 3473, control: 3026 mean RPKM), but this overall effect was not statistically significant (p = 0.36). ARGs conferring resistance to tetracycline (class level) were most strongly case-associated (case: 788, control: 620 RPKM, p = 0.01), especially within farms ES2 and ES3. Specifically, tetracycline and doxycycline resistance, which are largely conferred by the same ARGs, were the most abundant Resfinder ARG phenotype categories and had the strongest case-associations (Figure 4). *Tet(Q)* was the individual gene with the strongest case-association (case: 134, control: 75 RPKM, p < 0.01). ARGs of classes aminoglycoside, beta-lactam, and macrolide, which like tetracyclines are commonly used to treat *S. suis* disease, also trended towards higher abundance in case piglets (Table S4).

We examined the presence of case-associated ARGs in MAGs to determine whether the higher ARG abundance might be linked to high (chromosomal) ARG content in the case-associated taxa. We found that *tet(Q)* was only present in case-associated *Prevotella* MAGs, and case-associated ARGs *ant(6)-Ib* and *tet(44)* were only found in case-associated *Fusobacteriales* MAGs. ARGs found in control-associated *Rothia* and *Clostridium* MAGs, *lnu(P), erm(Q), tetA(P)*, and *erm(X)*, were not case-associated.

**Figure 4:**
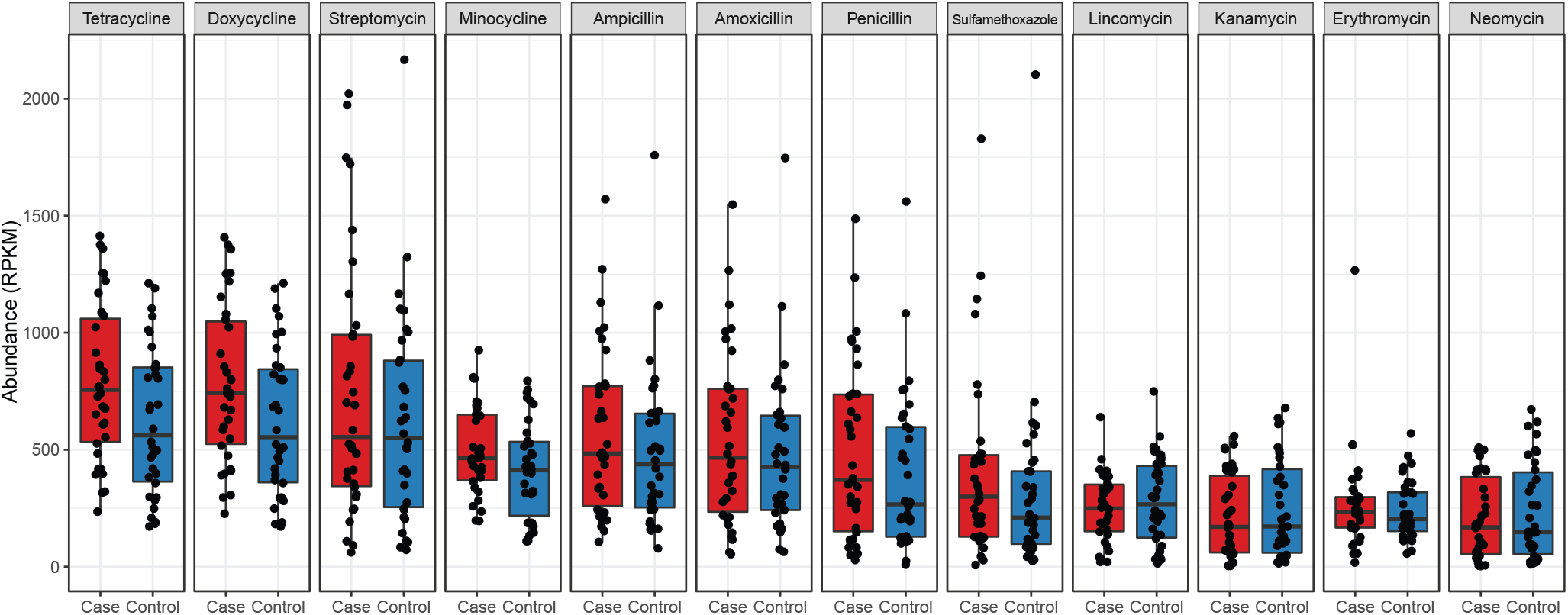
Total abundance of the 12 most abundant Resfinder ARG phenotype categories in case and control piglets. Note that some genes confer resistance to several different antimicrobials, making the sum of phenotype abundances greater than the total ARG abundance. For example, resistance to tetracycline and doxycycline is largely conferred by the same genes.

### Comparison of the tonsillar and nasal microbiota

To evaluate whether differences in microbiota composition between case and control piglets were specific to the tonsillar microbiota or a general trend in the upper respiratory tract, we collected 41 nasal swabs from three Spanish farms, ES1-3. The microbiota composition of the nasal and tonsillar swabs was significantly different, but the farm explained more variation (R2 = 0.08, p < 0.001, Bray-Curtis PERMANOVA) than nasal/tonsillar sampling (R2 = 0.06, p < 0.001). The nasal microbiota was characterised by higher abundance of genera *Moraxella* (nose: 32%, tonsil: 16%) and *Bergeyella* (nose: 5%, tonsil: 1%) and lower abundance of most other genera.

Case and control piglets had a significantly different nasal microbiota (R2 = 0.07, p = 0.04, Bray-Curtis PERMANOVA). This effect size was comparable to the difference found in the tonsillar microbiota samples for the same subset of piglets (R2 = 0.07, p = 0.09). However, the ASVs that were differentially abundant in the nasal microbiota of case and control piglets were not the same ASVs that were differentially abundant in the tonsillar microbiota. Some ASVs had inverse case-control associations: ASV 21 (*F. gastrosuis*), ASV 29 (*Prevotella* sp.) and ASV 3 (*Alloprevotella* sp.) were case-associated in the tonsillar microbiota, both overall and within ES2 and ES3, but health-associated in the nasal microbiota.

### *S. suis* diversity in the tonsillar microbiota

Amplicon sequencing data showed higher *S. suis* abundance in control piglets compared to case piglets within pre-weaning outbreak farms (farms ES3 and ES4, case: 3.9%, control: 6.4%, p = 0.02). This effect was not significant on post-weaning outbreak farms (DE1, DE3-6, ES2, NL1, case: 4.4%, control: 5.1%, p = 0.27).

A total of 89 ASVs were classified as *S. suis*, and 52 of these were present in two or more tonsil samples, suggesting that they were unlikely to be sequencing artefacts. Farm piglets were colonised by an average of 7 *S. suis* ASVs, ranging between 2 and 16. The relative abundance of *Streptococcus suis* was similar in outbreak and control farms (Figure 5A). Comparison of the 89 *S. suis* ASVs with the 16S rRNA gene V3-V4 region of 2463 *S. suis* assemblies available on NCBI assembly showed that many ASVs were poorly or not at all represented by sequenced genomes. This likely relates to the fact that many clinical but few non-clinical strains have been sequenced.

Unfortunately, ASVs are not good markers for assessing *S. suis* strain diversity. The V3-V4 region of the 16S rRNA gene correlates poorly with whole-genome phylogeny, and both clinical and non-clinical strains share the same ASVs. Among all *S. suis* assemblies, 82% shared a 16S rRNA gene V3-V4 region identical to ASV 6, which comprised 30% of the *S. suis* relative abundance in the tonsillar microbiota. ASV 17, the second most abundant *S. suis* ASV, comprised 21% of all *S. suis* relative abundance in the microbiota but was found only in 0.08% of the 2463 assemblies. In addition to strain DE512T1 collected in the present study, ASV 17 was only found in two isolates recently sampled from diseased pigs in China (GCF_019793915.1 and GCF_019794525.1).

We used metagenomic shotgun sequencing data to assess the relative proportions of commensal and pathogenic *S. suis* in the tonsillar microbiota of each piglet. First, we created a *S. suis* pangenome by clustering protein-coding genes from a previously published genome collection [24] at 80% identity, and mapped metagenomic reads to the representative sequences of each cluster (i.e., gene) to assess their abundance in each sample. We then calculated the ratio of prevalence of each cluster in clinical and non-clinical genomes to assess their putative association with pathogenicity. We found that in the tonsillar microbiota, genes predominantly found in non-clinical strains were more abundant than genes common in clinical isolates (Figure 5B). There was a small positive correlation between the ratio of gene clinical/non-clinical genome presence and ratio of case/control sample abundance (Figure 5C, Spearman R = 0.05, p < 0.01), due to higher abundance of commensal *S. suis* in control samples.

The gene *sly*, which encodes the pore-forming toxin suilysin, is commonly used as a genetic marker to distinguish clinical *S. suis* strains. *Sly* is well suited for use in metagenomic analysis due to high sequence conservation across *S. suis* strains, and it was found in the genome of at least 1 clinical isolate from all farms where we sequenced clinical isolates. The *sly* abundance was low in the tonsillar metagenomes, and similarly abundant in case and control piglets (Figure 5D). On some farms we did not detect *sly* in any sample, but this may be due to small sample size and insufficient sequencing depth.

**Figure 5:**
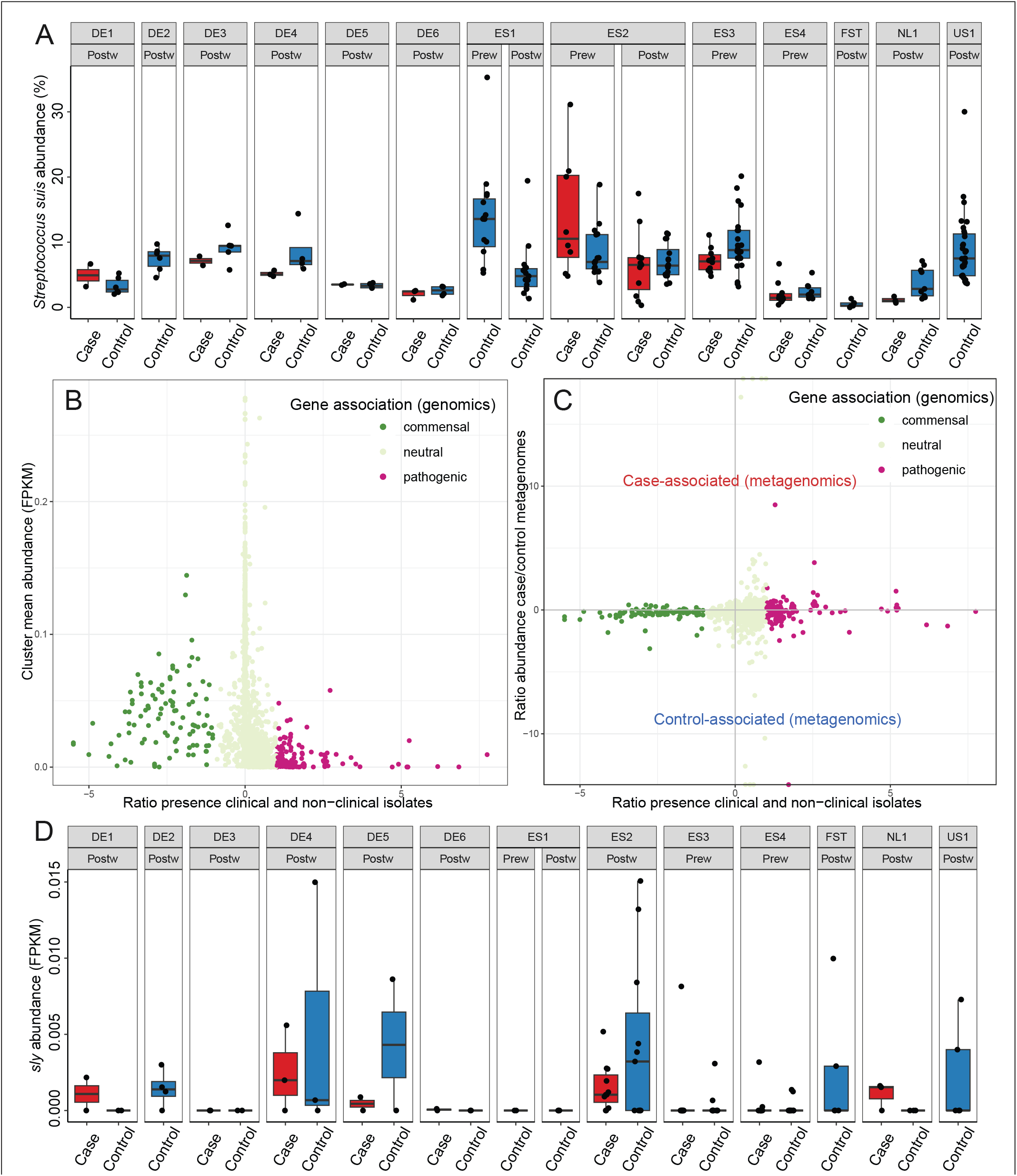
Abundance and diversity of *S. suis*. **A)** Total relative abundance of *S. suis* per farm (amplicon sequencing data). **B)** Within the *S. suis* pangenome, genes associated with non-clinical strains (x-axis) were more abundant (y-axis) in metagenomic tonsillar microbiota samples. Genes with a ratio of 0 were equally common in clinical and non-clinical strains, and genes with a ratio of 1 are twice as common in clinical strains. **C)** Correlation between gene association with clinical/non-clinical genomes (x-axis) and association with abundance in case-control metagenomic samples (y-axis). **D)** The abundance of the suilysin encoding gene *sly*, a gene highly conserved in clinical strains, in case and control samples per farm.

## Discussion

In this study, we found that the tonsil microbiota composition of case piglets with clinical *S. suis* signs was significantly different from the microbiota composition of asymptomatic controls. *Streptococcus suis* disease may occur as a part of polymicrobial infections collectively known as the porcine respiratory disease complex (PRDC), which includes porcine reproductive and respiratory syndrome (PRRS) virus, swine influenza A and potentially bacterial primary and secondary (opportunistic) pathogens [21, 32]. We found no association between the abundance of the species linked to PRDC and piglet case/control status in piglets with weaning-age systemic *S. suis* disease, but colonization by PRDC-associated taxa may be more relevant to respiratory disease in older finisher pigs.

We identified previously unreported disease-associations with *Fusobacterium gastrosuis, Bacteroides heparinolyticus*, and uncultured *Prevotella* and *Alloprevotella* species. *Fusobacterium gastrosuis* has previously been linked to *Helicobacter suis* infection in the gastric microbiota and shown to have genes involved in adhesion, invasion and induction of cell death, and immune evasion in other *Fusobacterium* species [33]. The uncultured case-associated *Alloprevotella*/*Prevotella*/*Bacteroides* species lack isolate genomes and are unknown in relation to *S. suis* disease, possibly because they are ‘unculturable’. Case-associated taxa may also interact with the host to facilitate *S. suis* crossing epithelial barriers without entering the bloodstream themselves, thus remaining undetected by necropsy. Alternatively, disease-associated taxa may increase in abundance due to host immune status and dysbiosis, as has been suggested for oral *Prevotella*/*Alloprevotella* species in humans [34].

Case piglets had lower tonsillar *S. suis* abundance than control piglets. This was due to reduced abundance of commensal *S. suis* clades, most of which are poorly represented among sequenced *S. suis* strains. While *S. suis* genes predominantly found in non-clinical isolates (i.e., putatively commensal-associated genes) were more abundant in control piglets, genes predominantly found in clinical strains, such as *sly*, were low in abundance and more equally distributed between cases and controls. This suggests that the majority of *S. suis* colonising tonsils are commensal, but also that disease-associated clades may colonise both symptomatic and asymptomatic piglets at low abundance. Based on these results, we conclude that tonsillar colonisation by *S. suis* itself cannot be used to reliably predict or even confirm ongoing *S. suis* invasive disease.

Associations between *S. suis* disease and the tonsillar microbiota have recently been investigated in North American piglets using amplicon sequencing by Niazy et al. 2022 [18]. Their study reported higher abundance of *Bacteroides* and *Lachnospiraceae* in controls, and higher abundance of *Campylobacter* and *Porphyromonas* in cases. Differences between studies may be explained by methodological differences and/or by European and North American piglets having fundamentally different microbiomes. We included one US farm in our study and found that these piglets had low microbiota diversity. Although most microbiota members were shared at the ASV level, composition was dominated by high abundance of *Actinobacillus* and *Streptococcus*. The different findings in Niazy et al. may also be due to them sampling whole tonsillar tissue, whereas we used swabs. In addition, they sampled piglets suffering from other pathologies such as rectal prolapse and hernia as controls, so the case piglets were not compared with healthy controls as in the present study. While their study may include control-associated taxa that are in reality associated with other diseases, our results may not be specific only to *S. suis* disease but include microbiome traits associated with low health status in general. Niazy et al. also sampled uneven numbers of case and control piglets per farm, and from some farms no controls. This confoundment of case-control and farm comparison may also have affected the results.

We found a trend for case piglets having a higher abundance of ARGs than control piglets, despite case-control pairs having received equal antimicrobial treatment. A similar effect of disease-associated resistome expansion can be observed in the human gut microbiome [35]. In the present study, ARGs conferring resistance to doxycycline and tetracycline had the strongest case-association, and ARGs conferring resistance to other antimicrobials used against *S. suis* also trended towards being more abundant in case piglets. ARGs found in MAGs of case-associated taxa *Fusobacterium, Prevotella*, and *Alloprevotella* had strong case-associations compared to those found in control-associated *Rothia* and *Clostridium* MAGs.

Although porcine *Rothia* isolates have high prevalence of ARGs [36], it is possible that more frequent antimicrobial exposure has exerted stronger selection pressure on disease-associated taxa, leading to accumulation of more ARGs.

While antimicrobial treatment may have exerted strong selective pressure over time, we found limited increase in the abundance of ARGs conferring resistance to the antimicrobials used to treat the sampled piglets. Antimicrobial use starting in the first few days after birth has been shown to influence composition and diversity in the piglet nasal microbiota [37], but in the present study most antimicrobial treatment started later in life, and only a few days before sampling. A second possible explanation for the limited effect of antimicrobial treatment on the tonsillar microbiota in our study may be the intrinsic resistance of biofilms to antibiotics [38]. The route of antimicrobial administration has been shown to influence the impact on the gut microbiota [39, 40], but it is not known to what extent antimicrobials provided in water, feed, or by injection are able to penetrate oral biofilm.

Farms had significant differences in microbiota composition, and piglets shared a similar microbiota composition with piglets within the same farm and country. This may be due to both sampler bias (a single person collected all samples in each country) and differences in farm environment, practices, and regulations. The piglet microbiota may for instance be influenced by factors such as cleaning, feed composition, temperature and antimicrobial treatment [41–48]. However, farms with comparable management practices did not show a more similar microbiota composition. Other relevant determinants of piglet microbiota composition may include host genetics and source of sows (including their vertically transferred microbiota). In both Spain and Germany, farms were spread over a large geographical area and, with the exception of ES1 and ES2, managed by different companies. Among the Spanish farms, farms ES1 and ES2, which were operated by the same company and had frequent exchange of animals, had the most similar microbiota composition. However, unrelated German farms shared a more similar microbiota composition than any of the Spanish farms.

We sampled the tonsillar microbiota of five piglets living outdoors in a forest in the Netherlands. These piglets had no recorded problems with diseases commonly affecting pigs on intensive farms, including *S. suis-*associated disease. We found that the tonsillar microbiota of forest piglets was fundamentally different from that of farm piglets and had lower ARG abundance. Many core ASVs in farm piglets were absent in the forest piglets, and core ASVs from the forest piglets were not found in farm piglets. The ecological farm DE4, where the piglets had straw bedding and outside access, did not have a more similar microbiota to the forest piglets than the conventional farms. Living outdoors may shape the tonsillar microbiota via exposure to environmental microbes from soil and diverse natural feed sources, and via high air quality and lower exposure to bacterial transfer from other piglets. Previous studies have shown differences in the faecal [49–51] and nasal [52] microbiota of wild pigs, but we are not aware of any studies on specific factors that may drive differences between the natural and domestic microbiota or between farms with varying disease problems. Understanding these factors may be key to preventing disease by opportunistic pathogens in pig farming.

To investigate whether early-life microbiota composition can predict disease incidence, we conducted longitudinal sampling on farm ES2, collecting prospective samples pre-weaning, prior to clinical sign development (post-weaning). Future case piglets had reduced alpha diversity compared to asymptomatic controls and piglets from control farm ES1. Asymptomatic siblings of case piglets had alpha diversity intermediate between symptomatic siblings and control piglets from litters without any *S. suis* cases. *S. suis* disease cases were concentrated in a limited number of litters, despite most of the herd remaining unaffected. This suggests that maternal effects involving early life immunity- or microbiota may be important in predisposing to *S. suis* disease and potential dysbiosis. Maternal effects may involve vertical transmission of a disease-associated microbiota at birth, but also differences in maternal immunity and premature depletion of antibodies [53]. *Streptococcus suis* disease most commonly occurs around the time colostral antibodies to *S. suis* start becoming depleted. Several studies have reported that in general, colostral antibodies vary highly in abundance and half-life between individual piglets and litters of different sows [54–58]. It is therefore possible that during *S. suis* outbreaks, some piglets may have sufficiently high levels of maternal antibodies to opsonise invading *S. suis*, while other piglets may lack sufficient levels of maternal immunity.

In conclusion, we found small but significant differences between the tonsillar microbiota composition of *S. suis* case piglets and asymptomatic controls. We discovered novel taxa associated with *S. suis* disease, and that *S. suis* abundance was higher in controls. The microbiota differences may originate from dysbiosis during early life prior to disease onset, but further research is needed to assert this. Case piglets had higher abundance of ARGs conferring resistance against antibiotics commonly used to treat *S. suis* disease. This may be linked to high ARG prevalence in case-associated taxa, which may face strong selective pressure for resistance due to more frequent exposure to antimicrobial treatment than control-associated taxa.

## Materials and Methods Animals

We utilized a case-control study design to assess the association between tonsillar microbiota composition and incidence of invasive *S. suis* disease. To search for consistent associations between microbiota composition and *S. suis* disease incidence, we included farms from different countries with different livestock management systems (Supplementary text 1). We obtained tonsil swabs from 3-to 10-week-old piglets at 13 farms, 9 of which had ongoing *S. suis* disease outbreaks. Three sampled farms had no history of *S. suis* disease. We compared our European farms with samples obtained at one US farm that had a history of *S. suis* disease but no cases at the time of sampling. Finally, we sampled 5 piglets free-living in a forest in the Netherlands at approximately 1 month after separation from the sows.

We selected 45 pairs of case-control piglets for metagenomic sequencing (Table 1). Pairs were selected to match as well as possible, coming from the same pen, room, and/or sow, and having received equivalent antimicrobial treatment. The cross-sectional case-control study design was extended with longitudinal sampling on two farms, ES1 and ES2, to determine whether microbiota differences in early life may be correlated with, or even predict, future *S. suis* disease occurrence. ES1 is a production farm free of *S. suis* disease, despite all sows and their carried microbiota and *S. suis* strains originating from ES2, which had severe *S. suis* disease problems.

**Table 1:** The number of case and control samples collected from each farm. See Supplementary text 1 for detailed information.

| Farm | Country | Age | 16S rRNA gene amplicon |  | Metagenomics |  |
| --- | --- | --- | --- | --- | --- | --- |
|  |  |  | Case | Control | Case | Control |
| DE1 | Germany | Postweaning | 2 | 6 | 2 | 2 |
| DE2 | Germany | Postweaning |  | 6 |  | 4 |
| DE3 | Germany | Postweaning | 2 | 5 | 2 | 2 |
| DE4 | Germany | Postweaning | 3 | 4 | 3 | 3 |
| DE5 | Germany | Postweaning | 2 | 4 | 2 | 2 |
| DE6 | Germany | Postweaning | 3 | 4 | 2 | 2 |
| ES1 | Spain | Postweaning |  | 15 |  | 3 |
| ES1 | Spain | Prewaning |  | 15 |  | 2 |
| ES2 | Spain | Postweaning | 11 | 16 | 11 | 11 |
| ES2 | Spain | Prewaning | 8 prospective | 16 |  |  |
| ES3 | Spain | Prewaning | 10 | 22 | 10 | 10 |
| ES4 | Spain | Prewaning | 10 | 10 | 10 | 10 |
| FST | Forest | Postweaning |  | 5 |  | 5 |
| NL1 | Netherlands | Postweaning | 3 | 9 | 3 | 3 |
| US1 | USA | Postweaning |  | 30 |  | 5 |

Clinical signs consistent with *S. suis* infection were recorded at each farm visit. Cases fell into two categories: arthritis (typically presenting as lameness) and meningitis (including otitis and sepsis, typically presenting as loss of balance, paralysis, paddling, shaking and convulsions). Confirmation of *S. suis* infection requires euthanasia and necropsy and was only carried out on a small subset of piglets due to welfare reasons. Most of the sampled piglets recovered after antimicrobial treatment. *Streptococcus suis* disease was confirmed by necropsy and isolation of *S. suis* from systemic sites in all 3 case piglets on farm NL1.

This study and all animal procedures were approved by the appropriate ethical committees. Sampling of the forest piglets was conducted according to the restrictions of the Animal and Human Welfare Codes in The Netherlands (2019.W-0026.001). On other farms, sampling was carried out for diagnostic purposes (covered by EU Directive 2010/63/EU).

### Whole genome sequencing of bacterial isolates

We selected 9 clinical (isolated from lesions observed at necropsy on the farms, but not necessarily from the same piglets sampled for tonsillar microbiota) and 7 non-clinical (from tonsillar swabs) *S. suis* strains for whole-genome sequencing. The isolates were grown in Todd-Hewitt broth with yeast extract overnight, and DNA was isolated using the PowerSoil DNA Isolation Kit (Qiagen) with 0.1mm silica bead beating. Isolated DNA was paired-end Illumina sequenced. Reads were trimmed with trimmomatic v0.39 [59], assembled with Spades v3.14.1 [60] and annotated using Prokka v1.14.5 [61]. Strain metadata and assembly statistics are shown in Table S5.

### Sample collection

The palatine tonsil microbiota of piglets was sampled by gently scraping the tonsillar surface with HydraFlock swabs (Puritan, ME, USA) for 10 seconds. The swabs were immediately placed in vials containing Powerbead solution (Qiagen, the Netherlands) and transported at -20 °C before being stored at -80 °C. DNA was isolated using the PowerSoil DNA Isolation Kit with 0.7mm garnet bead beating according to the manufacturer’s recommended protocol.

### Amplicon sequencing

The V3-V4 region of the 16S rRNA gene was amplified with primers 341F (5’-CCTAYGGGRBGCASCAG-3’) and 806R (5’-GGACTACNNGGGTATCTAAT-3’) and paired-end 250 bp sequenced using either the Illumina HiSeq 2500 or Novaseq 6000 platform. Reads were trimmed using cutadapt 2.3 [62] with default settings before being processed in DADA2 [29] following the v1.4 workflow for paired-end big data. Different sequencing batches were run separately before merging to account for differences in the learnErrors step. Taxonomy was assigned using SILVA database v138 [63]. Genus level taxonomy was assigned by the DADA2 pipeline using the RDP Naive Bayesian Classifier algorithm, and we used mmseqs2 easy-search with default settings to detect the species in SILVA with the highest identity alignment to each amplicon sequence variant (ASV) [64]. The species with the highest identity alignment above 98.5% was assigned. When several species aligned at equally high identity, all respective taxon names were assigned separated by a slash. ASVs with taxonomic assignment as eukaryote, mitochondrion, or chloroplast were discarded. Alpha and beta diversity were calculated on rarefied data (24325 reads) using R packages Phyloseq [65] and vegan [66]. The vegan::adonis function was used to perform PERMANOVA to determine the overall compositional differences between groups. Vegan function RDA was used for principal component analysis (PCA) and redundancy analysis (RDA).

### Shotgun sequencing

Metagenomic libraries were prepared using the NEB Next Ultra DNA Library Prep Kit (New England Biolabs, ME, USA) following the manufacturer’s instructions. DNA was fragmented to 350 bp, purified using AMPure XP (Beckman Coulter, CA, USA) and sequenced with 150 bp paired-end sequencing on an Illumina NovaSeq 6000 machine.

### Analysis of metagenomic data

Pig and plant (feed) reads were removed with kneaddata (https://github.com/biobakery/kneaddata) using the genomes of pig (GCF_000003025.6), wheat (GCF_002162155.1) and maize (GCF_902167145.1). As some samples still had a high proportion of host or plant reads left after this step, we further normalised read counts by the proportion of plant and pig reads found by Kraken analysis [67]. This prevented samples with low proportions of bacterial reads from being outliers when mapping metagenomic reads to marker genes and metagenome assembled genomes (MAGs). We generated MAGs using MetaWRAP v1.3.2 [68] with SPAdes v3.14.1 [60], and identified MAG taxonomy using GTDBtk v1.3.0 [69] and ARGs with ResFinder software v4.1 [31]. We also used the ResFinder database to quantify ARG abundance directly from metagenomic reads. Before mapping, we clustered the genes in the database to 90% identity using mmseqs2 [64] easy-cluster with settings ‘--min-seq-id 0.9 --cov-mode 0’. Reads were mapped to the representative sequences of the clusters using mmseqs2 easy-search with default settings, and reads aligning with minimum 100 bp and 95% identity were accepted.

To assess the *S. suis* population in the tonsillar microbiota, we created a *S. suis* pangenome by clustering all protein-coding genes and mapping metagenomic reads to representative sequences. We used previously published metadata covering 1703 assemblies [24]. We annotated the genomes using Prokka v1.14.5 and clustered all protein coding genes at 80% residue identity using mmseqs easy-cluster (-min-seq-id 0.8 -cluster-mode 2 -cov-mode 1). We determined the association of each cluster with presence in clinical and non-clinical strains by calculating the ratio of percentage presence in each group. The ratio of clusters more prevalent in clinical strains was calculated as (% presence clinical/% presence non-clinical) - 1, and clusters more prevalent in non-clinical strains by ((% presence non-clinical/% presence clinical) * -1) + 1, such that clusters equally prevalent in clinical and non-clinical strains had a ratio of 0. We mapped metagenomic reads to the representative sequence of each cluster as described above and accepted reads mapping at 80% identity and 80% length.

## Data availability

All microbiome sequencing data generated in this study have been deposited in the NCBI BioProject portal under accession number PRJNA854341. Genome assemblies are available under accession numbers PRJNA849547 and PRJNA849577.

## Acknowledgements

We thank the farmers and veterinarians who participated in the study. Most farms are anonymised. Farm NL1 is Schothorst Feed Research B.V. (www.schothorst.nl) and the forest piglets (FST) were collected from Boeren in het Bos (www.boereninhetbos.nl).

