## Supplementary text for "Streptococcus suis infection on European farms is associated with an altered tonsil microbiome and resistome"

**Additional file 1: Supplementary information - Information on farms.**

This document contains information on farms relevant to understanding the farm practice and disease situation. The information provided is qualitative and based on information from veterinarians and farm workers. The study farms were selected based on previous contact and represent a wide range of *S. suis* problem scenarios and are not intended to be representative of the different countries. The Spanish farms were sampled during severe outbreaks with a subset of symptomatic piglets sampled, while German and Dutch farms were sampled during smaller outbreaks with all available case animals being sampled.

**Table 1.** Farm overview.

| **Farm** | **Country** | **Symptoms** | **Disease stage** | **Farm disease history** |
| --- | --- | --- | --- | --- |
| DE1 | Germany | Meningitis | Post-weaning | Yes |
| DE2 | Germany | None | None | None |
| DE3 | Germany | Meningitis | Post-weaning | Yes |
| DE4 | Germany | Meningitis | Post-weaning | Yes |
| DE5 | Germany | Meningitis | Post-weaning | Yes |
| DE6 | Germany | Meningitis+Arthritis | Post-weaning | Yes |
| ES1 | Spain | None | None | None |
| ES2 | Spain | Meningitis+Arthritis | Post-weaning | Yes |
| ES3 | Spain | Arthritis | Pre-weaning | Yes |
| ES4 | Spain | Arthritis | Pre-weaning | Yes |
| NL1 | Netherlands | Meningitis | Post-weaning | Yes |
| NL2 | Netherlands | None | None | None |
| US1 | USA | None | None | Yes |

**Table 2.** Antimicrobial administration in farms.

| **Class** | **Type used in study piglets:** |
| --- | --- |
| aminoglycoside | Neomycin (ES3), Gentamicin (ES4, US1) |
| tetracycline | Tetracycline (DE1, DE3) |
| beta_lactam | Amoxicillin (DE3, DE6, ES2, ES3, ES4, US1) |
| fluoroquinolone | Marbofloxacin (ES2) |

**Fam description:**

**DE1 (Germany)**

Regular commercial farm. Intermittent *S. suis* problems. Tetracycline treatment since day 1 for piglet 1-4 but not 5-8. Farm size: Sows: 800, Nursery: 2000.

**DE2 (Germany)**

Regular commercial farm. Very high health status, and piglets had no problems with any disease in the period around sampling. Prestarter feed provided in both dry and liquid form and open water source provided. The piglet did not have their teeth ground or cut. All piglets were treated with amoxicillin the first day after birth as is typical in German farms. Farm size: Sows: 500, Nursery: 1200.

**DE3 (Germany)**

Regular commercial farm with some PRRSV problems (no vaccination) and intermittent *S. suis* problems. Low health farm with high antimicrobial usage for months/years. Amoxicillin and Tetracycline treatment for 5 days before sampling of piglets 1-5 but not piglet 6-7. Farm size: Sows: 600, Nursery: 2800.

**DE4 (Germany)**

Ecological farm utilizing vaccines. The piglets have outside access and straw bedding. The farm has reduced biosecurity, not only due to the outside access and lower level of cleanliness but also due to a lack of separation of airflow between age groups, which may facilitate transfer of pathogens. In addition to *S. suis*, the piglets have a high burden of parasites as well as rotavirus problems. No antimicrobial treatment of the sampled piglets. Farm size: Sows: 300, Nursery: 1200.

**DE5 (Germany)**

Regular commercial farm. Intermittent *S. suis* problems. No antimicrobial treatment of the sampled piglets. Farm size: Sows: 1200, Nursery: 5000.

**DE6 (Germany)**

Regular commercial farm. This farm has a history of severe *S. suis* problems. PRRSV and PCV2 (outbreak ongoing at sampling, may relate to failure in vaccination), and other pathogens also cause problems at this farm. Only 1 pig treated with antimicrobials (Amoxicillin) in the days prior to sampling. Farm size: Sows: 100, Nursery: 2000.

**ES1 (Spain) - sample prefix QT**

Regular commercial farm. Production farm receiving sows from ES2. No history of *S. suis* problems, although this is not a high health status farm in general. No antimicrobials used on sampled piglets. This farm was visited two times, with microbiota sampling of the same piglets at day 14 (1 week before weaning) and day 51 (3 weeks post-weaning). Weaning at day 21.

**ES2 (Spain) - sample prefix RC**

Regular commercial breeding farm. This farm delivers dams to farm ES1. This farm has a history of severe *S. suis* problems. This farm was followed from 1 week before weaning to 3 weeks post-weaning, with 1 microbiota sampling per week for 5 weeks. Weaning day 26-29. Sampling at day 18-21 (week -1), day 24-27 (week 0), day 32-35 (week +1), day 39-42 (week +2), and day 47-50 (week +3). All piglets were given amoxicillin in water at day 39, 12 days after weaning and 1 day before sampling point week 2 after weaning. Piglets in some nursery pens were treated with intramuscular marbofloxacin injections, see metadata.

**ES3 (Spain) - sample prefix BN**

Regular commercial farm. We sampled pre-weaning *S. suis* cases. This farm has a history of severe *S. suis* problems. All piglets were given prestarter feed with amoxicillin and neomycin.

**ES4 (Spain) - sample prefix JT**

Regular commercial farm. This farm has a history of severe *S. suis* problems. All piglets were given amoxicillin + gentamicin injections 2 days before sampling.

**NL1 (Netherlands)**

Research farm, Schothorst Feed Research B.V., Lelystad, Netherlands. High health status and low antimicrobial usage. Intermittent *S. suis* problems. No antimicrobials were given to the sampled piglets.

**FST (Netherlands)**

Piglets free roaming in a forest, with no access to indoor housing. Breed: Tamworth. No vaccination or antimicrobial treatment. The piglets forage for food, but this is also supplemented with some dry feed, fruit, and vegetables. Piglets were weaned and living in a different section than the sows at the time of sampling but spent considerably longer time with the sow than in regular farms (until naturally weaned). The outdoor conditions, rooting in soil, varied diet, gradual weaning, and rare breed may all have contributed to the diverged tonsillar microbiota.

**US1 (USA)**

Regular commercial farm. A farrow-to-finish farm located in the Midwest USA. Overall high health status. Intermittent *S. suis* problems, usually occurring late nursery or early finisher stage. All piglets were treated with gentamicin (day 26) and amoxicillin (day 45). The piglets were weaned at day 26 and sampled at day 55/62.
