## Supplementary figures for "Streptococcus suis infection on European farms is associated with an altered tonsil microbiome and resistome"

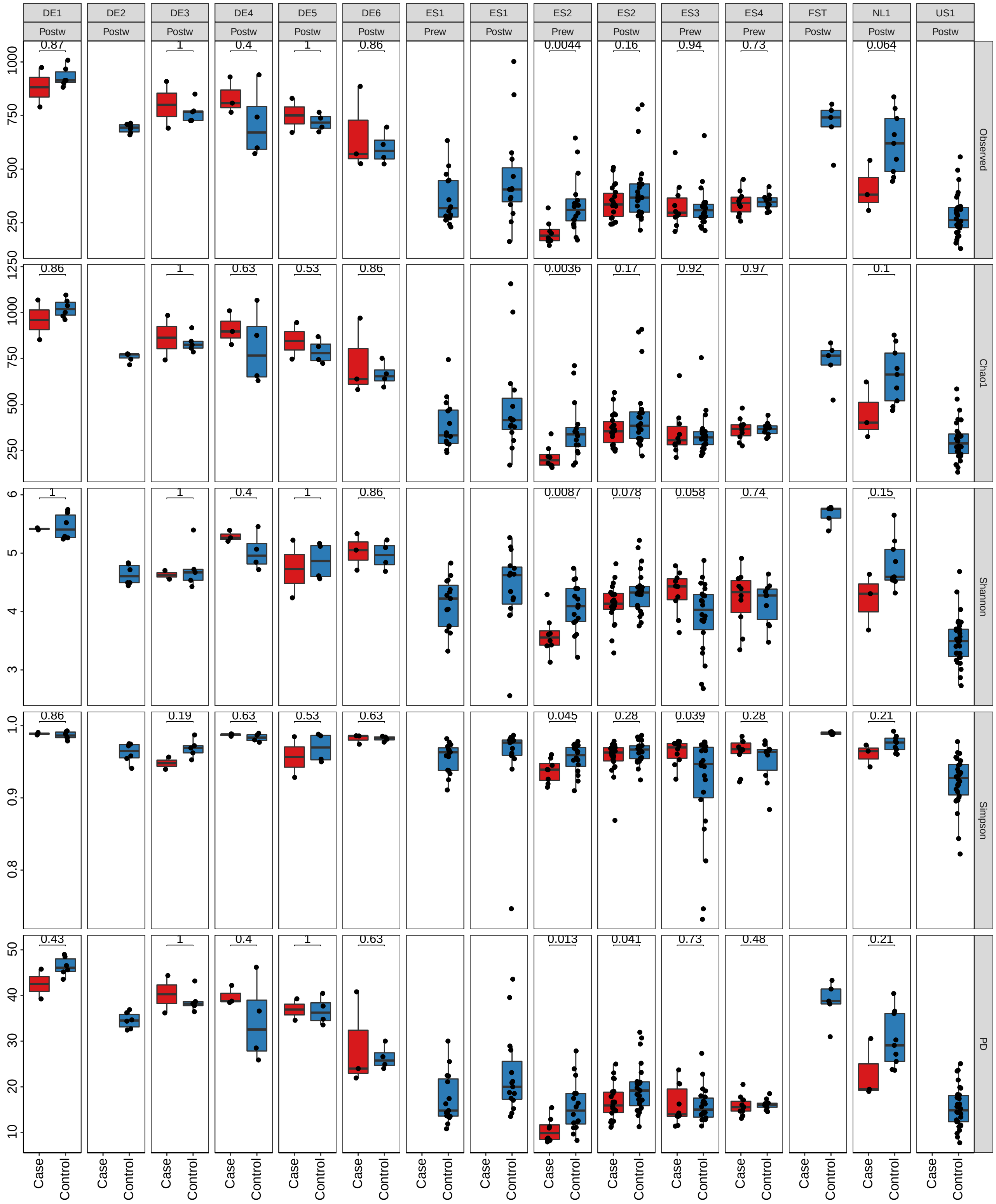


**Figure S1:** Comparison of alpha diversity in case and control piglets within each farm and age group. Plots based on different diversity metrics are shown facet on the y axis. The figure is based on amplicon sequencing data. Observed = the total number of ASVS detected in each sample. PD = phylogenetic diversity.
